# HD-tDCS over mIPS causally modulates online reach correction

**DOI:** 10.1101/708693

**Authors:** Alexander Goettker, Brandon Caie, Jerrold Jeyachandra, Sisi Xu, Jason Gallivan, Jacek Dmochowski, Katja Fiehler, Gunnar Blohm

**Affiliations:** Justus-Liebig University Giessen, 35394 Giessen, Germany; Center of Neuroscience Studies, Queen’s University, Kingston, ON K7L 3N6, Canada; Department of Biomedical Engineering, City College of New York, New York NY 1003; Center for Mind, Brain and Behavior (CMBB), Universities Marburg and Giessen, Germany

**Keywords:** Motor control, arm movements, parietal cortex, transcranial electrical stimulation, EEG

## Abstract

Brain lesion and stimulation studies have suggested posterior parietal cortex and the medial intraparietal sulcus in particular as a crucial hub for online movement error corrections. However, causal evidence for this is still sparse. Indeed, lesion studies are potentially confounded by compensatory reorganization mechanisms while brain stimulation studies have produced heterogeneous results when employing transcranial magnetic stimulation. Here we designed a new complementary paradigm using fMRI-guided high-definition transcranial direct current stimulation (HD-tDCS) of the left medial intraparietal sulcus (mIPS) together with regression-based mediation analysis to re-examine the causal role of mIPS in online reach corrections to jumping targets. We obtained two independent measures of stimulation-induced changes in brain activity by modeling current flow in the brain and through EEG recordings before and after HD-tDCS stimulation. Third, to quantify behavioral effects of HD-tDCS we computed movement curvature as a measure of online correction. We demonstrate that both of our measurements of brain activity were consistent with a polarity-specific modulation of the online correction for targets jumping to the contralateral side of the stimulation. Importantly, using a mediation analysis of the relationship between stimulation current and movement curvature suggests that the induced current modifies brain activity, which then leads to the observed behavioral changes. This unique combination of methods and analysis thus provides complementary evidence for the crucial role of the posterior parietal cortex in online error correction, while at the same time setting a new methodological standard with respect to the causal influence of transcranial direct current stimulation.

**New & Noteworthy:** Transcranial direct current stimulation (tDCS) is an interesting and potentially useful tool for asking causal scientific questions and design clinical treatments. With our unique combination of highly accurate fMRI guided stimulation, current forward modeling, EEG recordings before and after the stimulation and behavioral changes we could unravel the causal structure of tDCS. Our approach naturally deals with the variability of tDCS results, increasing its potential usefulness as a tool for research and clinical applications.

## Introduction

In order to interact with objects in our environment we often bring our hands to the object to manually explore and manipulate them. Those movements can be inaccurate due to sensory-motor noise (Harris & Wolpert, 1998; van Beers, Baraduc & Wolpert, 2002; Faisal, Selen & Wolpert, 2008) or sudden changes in the environment. To compensate for errors, anonline-control system (see Pelisson, Prablanc, Goodale & Jeannerod, 1986; Pisella et al., 2000; Prablanc, Desmurget & Grea, 2003; Gaveau et al. 2014) compares an continuously updated representation of the target position and an internal estimate of the hand state (Prablanc & Martin, 1992; Desmurget et al. 2001). While the target position is usually determined by visual information (Sarlegna et al., 2003), the internal estimate of the hand state relies on a combination of somatosensory feedback and efferent information of the motor command (Miall & Wolpert, 1996; Scott, 2012; Sarlegna et al., 2015). Multiple lines of research point to the posterior parietal cortex (PPC) as one of the important neural structures for online control, as evidenced by fMRI (Ogawa et al. 2006; Limanowski et al., 2017), TMS (Desmurget et al.1999; Tunik et al. 2005) and lesion studies (Pisella et al.,2000; Grea et al.,2002; Rossetti et al., 2003; Hwang & Anderson, 2012; Andersen et al. 2014).

Despite this accumulation of evidence, not much is known about the underlying neural mechanisms during online control (see Battaglia-Mayer et al. 2014; Archambault et el., 2015).In addition, a recent TMS study was not able to replicate an effect on online control of hand or foot movements (Marigold et al., 2019), suggesting a broad network of areas with varying specializations in the PPC (Medendorp & Heed, 2019).Furthermore, lesion studies (see Cavina-Pratesi, Connolly & Milner, 2013; Buiatti, Skrap & Shallice, 2013) provide evidence for causal involvement of the PPC, but cannot point to the exact mechanisms underlying behavioral differences(Jonas & Kording, 2017) and suffer the confound of compensatory brain reorganization mechanisms (see Rorden & Karnath, 2004).

Therefore, we aimed at reinvestigating the causal role of the PPCin online control by using transcranial direct current stimulation (tDCS). In contrast to TMS, which disrupts the cortical output of an area through the generation or inhibition of spikes (Siebner, Hartwigsen, Kassuba, & Rothwell, 2009), tDCS is capable of bidirectionally influencing dendritic input through modulation of pre-synaptic activity (Bączyk & Jankowska, 2014, see Liu et al. 2018). One advantage of tDCS is that we can modelin duced current density (Dmochowski et al, 2011; Saturnio et al, 2015, 2018) to estimate its orientation and magnitude in the target area and predict changes in brain activity and inter-individual differences on the behavioral level. We targeted the middle intraparietal sulcus (mIPS), a region in the PPC, which is involved in movement planning and online control (Grea et al., 2003;Vesia et al., 2010; Gallivan et al., 2011; Gertz & Fiehler, 2015).

Thus far, previous research has looked at the direct relationship between tDCS and behavior (e.g. Jacobsen et al., 2012) or used EEG to independently measure and verify the electrophysiological changes induced by tDCS (e.g. Mangia et al, 2014; Lauro et al, 2014). However, to our knowledge our study is the first one that combines all these measures: we designed a mediation analysis (Baron & Kenny, 1986) based on the modeled current, changes in brain oscillations and changes in behavior to reconstruct the causal structure of the stimulation. By leveraging the double dissociation in current polarity afforded by tDCS and our unique set of measurements, we demonstrate that the induced current is related to behavioral changes, but importantly this relationship is mediated by altered brain activity. By applying such an interventional methodology (Marinescu, Lawlor & Kording, 2018), our results provide causal evidence for the role of mIPS in online reach correction, while at the same time allowing insight into relevant mechanisms.

## Methods

### Overview

By stimulating the left mIPS with HD-tDCS we aimed to provide complementary evidence for a causal role of the PPC during the online control of right-arm reaching. We combined detailed psychophysics, current forward modeling and independently measured stimulation-induced EEG changes to establish a causal relationship between the induced current, changes in brain activity and the altered behavior. We made use of the bidirectional effect of tDCS by stimulating all participants with anodal and cathodal stimulation in randomized order on two different testing days, separated by at least one week. To achieve anatomically accurate stimulation of the targeted area, we used task-based fMRI to localize the left mIPS in each participant and registered the location on the skull (see fMRI **localizer task** and **preparation of stimulation**) to place the electrodes for the stimulation accordingly (see **HD-tDCS**). Based on the placement of the electrodes and the participant’s brain scan we were able to model the current flow and quantify the expected effect of the stimulation at the targeted site on a subject-by-subject level (see **Current Forward Modeling**). For the behavioral experiment, each day was divided into three different consecutive sessions (baseline, stimulation, and post-stimulation). To independently measure changes in brain activity caused by the stimulation we used the same electrodes as during HD-tDCS stimulation to record the EEG response in the baseline and post-stimulation sessions (see **EEG Recording)**. During the baseline session participants performed four blocks without any stimulation to obtain a representative measurement of their performance. The task was to reach to the location of visually presented dots, and some of them jumped to a new location at hand movement onset (see **Experimental setup and paradigm**). During the stimulation session participants also completed four blocks, but now with constant direct current stimulation. The stimulation duration was 20 minutes, and all of the participants finished their four blocks during the time of stimulation. After stimulation, we measured post-stimulation trials, where participants again completed four blocks without stimulation. We quantified the online correction performance by measuring the curvature of the respective movements (see **Behavioral Data**). This combination of measurements allowed us to perform a mediation analysis (see **Mediation analysis** and **Simulating mediation statistics**) to establish a relationship between the induced current and behavioral changes, while showing that the behavioral changes are mediated by underlying changes in brain activity as measured through EEG.

### Participants

Ten healthy right-handed volunteers participated (age range: 20 – 39 years; 7 females) in the experiment. All participants had normal or corrected-to-normal vision. All participants gave informed written consent to procedures that were approved by the Queen’s University General Board of Ethics. Participants received financial compensation for their participation. One participant was excluded from the analysis as he did not exhibit online corrections during his baseline testing session.

### fMRI localizer task

To localize the left mIPS in each participant, all participants took part in a localizer testing session in an MRI-scanner prior to the main experiment (see figure 1B).MR imaging was performed at the Queen’s MRI facility using a 3-Tesla Magnetom Trio scanner (Siemens Medical Systems, Erlangen, Germany). The left mIPS was functionally localized on an individual basis using an interleaved center-out reaching task (Vesia et al., 2010; Vingerhoets, 2014).

**Figure 1.**
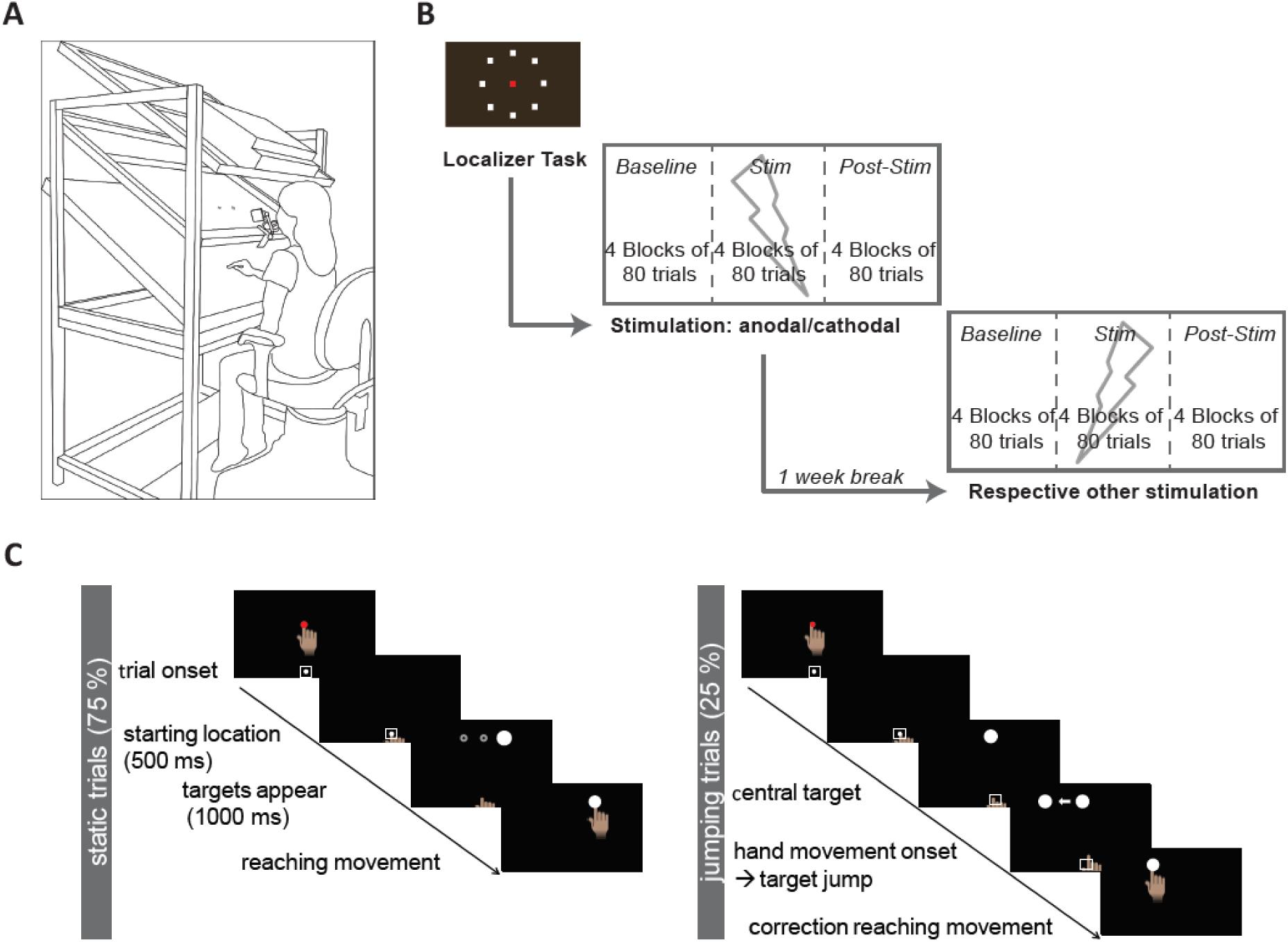
Overview of experiment. (**A**) Sketch of experimental setup. Participants sat in a chair in front of the tilted surface, which they viewed through a semi-silvered mirror. In this way, visual feedback about their hand position was occluded. (**B**) General overview of the different sessions. After they completed the functional localizer task in the MRI scanner, participants completed the main experiment on two testing days, separated by one week. (**C**). Depiction of the different trial types during the experiment. In static trials participants had to perform reaching movements to one of three possible target locations. In the jumping trials participants started their hand movement to the central target but it jumped either to the left or the right at hand movement onset.

Inside the MRI, the participant rested supine with a 32-channel head coil worn on the head. All stimuli were rear-projected using an LCD projector (NEC LT 265 DLP projector; resolution, 1024 × 768; 60 Hz refresh rate) onto a screen mounted behind the participant, and viewed by the participant via a mirror mounted to the head coil directly above the eyes. In each task block, the participants were presented with a circular arrangement of 8 white circles (∼15 degrees of visual angle) surrounding a central red or green square. When the central square was red, participants were instructed to make self-paced center-out reaching movements (small pivots about the wrist and elbow with their index finger) while fixating on the central fixation square; when the central square was green participants were instructed to make self-paced center-out saccadic eye movements while keeping their hand in a relaxed home position on their chest. The reach-saccade localizer included 8 reaching blocks (16 s per block), 8 saccade blocks (16 s per block), and 3 fixation/baseline blocks (10 s per block), which were placed at the beginning, middle and end of the run. The localizer run lasted a total of 5 min 22 s (157 brain volumes).

Functional MRI volumes based on the blood oxygenation level-dependent (BOLD) signal were collected using T2*-weighted gradient echo planar imaging with 240 mm x 240 mm field of view with 80 × 80 acquisition matrix size for an in-plane resolution of 3mm, repetition time (TR) = 2 s; time to echo (TE) = 30 ms, and flip angle (FA) = 90°. Each volume comprised of 35 contiguous axial slices (slice thickness 4mm) acquired in an interleaved order along the anterior commissure-posterior commissure (AC-PC) line. A 176 slice, high-resolution T1-weighted structural volume was acquired along the same orientation as the functional images using a 3D MP-RAGE sequence (single shot, ascending sequence in the sagittal plane with TR = 1760 ms, TE = 2.2 ms, FoV = 256 mm, flip angle = 90°, and voxel size = 1 mm^3^).

Data from the localizer run were registered to the corresponding structural image, aligned on the plane between the anterior commissure and posterior commissure. All preprocessing and univariate analyses were performed using Brain Voyager QX version 2.6 (Brain Innovation). Preprocessing for the localizer data included slice scan-time correction, 3D motion correction (alignment of the first volume of the functional scan, which was closest in time, to the anatomical scan), high-pass temporal filtering of 3 cycles/run, functional-to-anatomical co-registration, and spatial smoothing (trilinear-sinc interpolation performed during realignment, sinc interpolation performed during reorientation). We carried out subject-level analyses using a general linear model (GLM) with condition predictors created from boxcar functions convolved with a double-gamma hemodynamic response function (HRF). These were aligned to the onset of each reach and saccade block, with its duration based on block length. In a whole-brain analysis for each, the BOLD signal was contrasted between the reaching condition and the fixation condition to identify the hotspots of the left mIPS. The location of the left mIPS was defined by selecting the voxel peak medial to the intraparietal sulcus in the left hemisphere (Vingerhoets, 2014).

### Preparation of stimulation

During the main experiment, we used neuronavigation (Brainsight, Rogue Research Inc., Montreal, Quebec) to place the central electrode on the scalp over the fMRI-identified location of the left mIPS, so that the peak electric field would be directed to the mIPS. The participant-image registration used the landmarks of the nose tip, the nasion and the left and right intertragal notches. After the registration, the target location on the scalp was identified as the inline projection of the selected target region mapped onto the scalp that minimized the distance between target region and the target location on the skull.

After the registration of the target location on the scalp with Brainsight, a 10-20 system EEG mesh cap was fastened on the head of the participant. Embedded plastic holders in the mesh cap were used as holders for the electrodes. The central electrode was put directly above the target location on the scalp (by adjusting the cap), and additionally 4 pick-up electrodes were placed in a 3 cm radius surrounding the central electrode, leading to a 4×1 electrode configuration for HD-tDCS (Datta et al., 2009).

### Experimental setup and paradigm

Participants sat in a dark room at a tilted reaching setup (see figure 1A). Their head was immobilized by a personalized dental impression, attached to the setup. The tilted reaching setup consisted of an overhead monitor (ViewPixx 1920*1200 Pixel; 52 *29 cm, VPixx Technologies Inc., Saint-Bruno, QC Canada) for the presentation of the visual stimuli, a tilted reaching surface below the screen and a semi-silvered mirror positioned exactly in between the reach surface and the screen so that stimuli presented on the screen were projected onto the reach surface. The mirror allowed the participants to see the visual stimuli, but occluded the sight of their hand. All surfaces were tilted by 30°, which allowed for comfortable reaching and facilitated eye-tracking. The eye-tracker was attached to the frame of the semi silvered-mirror. The eye movements were tracked with an EyeLink 1000 eye tracking system (SR Research; sampling rate, 500 Hz; accuracy, 0.5°). Hand position during the reaching was recorded by localizing the 3D position of infrared light-emitting diodes taped to the tip of the right index finger (Optotrak Certus; Northern Digital; sampling rate 400 Hz; accuracy, 0.1 mm).

Participants were asked to perform reaching movements from an initial starting location to a visual target at one of three possible target locations. In some of the trials, the visual target changed position at the onset of hand movement. At the beginning of each trial, a white dot (30 pixels diameter) indicated the starting location. The starting location was presented on the horizontal midline slightly above the bottom end of the screen. Participants had to place their right hand at the starting location and were told to also look there. Although participants could not see their own hand in the experiment, during the beginning of the block or after the end of the previous trial a red marker indicated their hand position within 5cm of the start position to guide them back to the starting location. When the hand was kept at the starting location for 500 ms (defined as a frame of 50 pixels around the starting location) the starting location disappeared and a dot representing the target locations appeared. The central target was presented in a distance of 23 cm above the starting location, at the midline of the screen. The left and the right target locations were placed at the same height, but displaced laterally by 7.5 cm relative to the center target. As soon as the target appeared, participants were instructed to make a reaching movement to the target location and were allowed to move their eyes. The target stayed on for 1000 ms and afterwards the starting location appeared to signal the start of the next trial (see figure 1C).

Each block had a total of 80 trials, lasting approximately 4.5 minutes. Across trials the different target locations were randomized. Of the 80 trials, 20 trials changed the target position at hand movement onset. This small amount of target jump trials mitigated participants ability to predict jump trials. For the static targets the target was equally likely to appear at each of the target locations (left, center, right). The 20 trials jumping trials initially appeared at the central location, but jumped with equal probability to the left or right at hand movement (jumping left, jumping right). Hand movement onset was calculated online as the moment where the hand was outside of the initial frame around the starting location. In total, participants completed 1920 trials (2 polarities * 3 sessions * 4 blocks * 80 trials) including 480 jumping trials per participant.

### EEG Recording

During the baseline and post-stimulation session we used the central and four surrounding electrodes to record the EEG-response. This allowed us to assess the influence of HD-tDCS on oscillatory signatures of reaching. EEG data was acquired from the same electrodes delivering the stimulation using a 16 channel USB amplifier (V-AMP, Brain Products). Reference and ground electrodes were placed on the seventh cervical vertebrate and the olecranon respectively to minimize noise from muscular activity. EEG data was sampled at 2kHz and converted from analog to digital with a resolution of 24 bits at 0.0489 µV/bit. Event timing was tracked by a trigger shared between the EEG amplifier and the Eyelink 1000, allowing for behavioral events to be time-locked to the EEG signal.

### HD-tDCS

The electrodes were sintered Ag/AgCl ring electrodes (HD-target package, Soterix Medial Inc., New York, NY, USA) with a diameter of 1 cm. To improve the conductance, the hair was separated under the scalp using cotton swabs and the plastic holders were filled with conducting gel. An impedance check was performed prior to the experiment to ensure appropriate impedance limits (< 20 kΩ).

The stimulation was administered for 21 minutes. During the first 30 seconds the current strength ramped up and during the last 30 seconds ramped down again, leading to 20 minutes of constant stimulation with a current strength of 2mA. The stimulation was controlled by microprocessor-controlled constant current source (DC-STIMULATOR MC 8, NeuroConn GmbH, Ilmenau, Germany) to deliver continuous direct current. The direction of the current led to either anodal or cathodal stimulation under the central electrode.

## Data Analysis

### Behavioral Data

First, the data obtained by the Optotrak and Eyelink were low-pass filtered by an autoregressive forward backward filter with 50Hz cut-off frequency. Then, velocity and acceleration were computed using a central difference algorithm. The offline analysis was performed in MATLAB (The Mathworks, Natick, MA). The hand movement onset was defined as the point where the hand was moved outside of the initial frame around the start position. The end position was defined as the time when the velocity of the hand was lower than 1 cm/s.

The onset of online correction was defined by the interpolation method of Widjens et al. (2014).The average x-position of a movement to the central target was calculated for each session. Afterwards the difference between the x-position of each single jumping trial and the average for the central location was computed. This gave us a difference function, where we performed a linear regression between the points of 25% and 75% from the absolute maximum difference. The point where this line crosses zero was used as the onset of online correction. Based on the detected movement endpoints we computed the horizontal- and vertical-error and the precision of the movements. Horizontal- and vertical-error were based on the distance between the endpoint and the target location and the precision of the movements was defined as the area of an error ellipse which contained 95% of the movement endpoints.

The main variable that we used to quantify the online correction performance was the normalized curvature. The normalized curvature was defined as the maximum distance between the movement trajectory and a straight line which connected the start and the endpoint of the hand movement, divided by the length of the trajectory of the movement. Additionally, we did not only look at the absolute curvature of the trajectory, but also analyzed the development of curvature over time. Therefore, we aligned all trials on the calculated onset of the online correction and calculated the curvature in the same way with a sliding window of 100 ms in steps of 10 ms. We normalized the curvature over time based on the maximum curvature for the respective baseline session and then compared the peak curvature depending on the session and stimulation type. To quantify the effect of the stimulation on behavior we computed the difference between the stimulation and the baseline session for anodal and cathodal stimulation for our two measurements of curvature, absolute curvature and the peak curvature in the time course after correction onset. We collapsed the data over the four blocks of each session (except for the baseline where we only used the last 3 blocks, see exclusion of trials) and calculated the mean and the standard deviation for each participant for each of the sessions for the different variables. To test for systematic influences on the movement endpoints we ran a repeated measures ANOVA with the factors polarity (anodal, cathodal), session (baseline, stimulation, post-stimulation) and target type (static vs online corrected) separated for targets on the left and right. To test for effects of the stimulation on online correction we conducted repeated measures ANOVAs with the factors polarity (anodal, cathodal), target location (jumping left, jumping right) and session (baseline, stimulation, post-stimulation). As post-hoc analysis we conducted one-way ANOVAs with the factor session for either the anodal or cathodal values or post-hoc t-tests with Bonferroni correction.

### Current Forward Modeling

We simulated the induced electric field at the site of stimulation by constructing a tetrahedral mesh from the T1 and T2-weighted MRI of each participant and numerically solving a forward model for Maxwell’s equations across tissue boundaries (see Figure 2). In tDCS, the current flows through a heterogeneous medium with a complex spatial distribution that depends on both head geometry and the electrical conductivities of the various tissues. Thus, it is important to consider the influence of individual differences in anatomical properties such as cortical surface folding and skull thickness on the current path from scalp to target.

**Figure 2.**
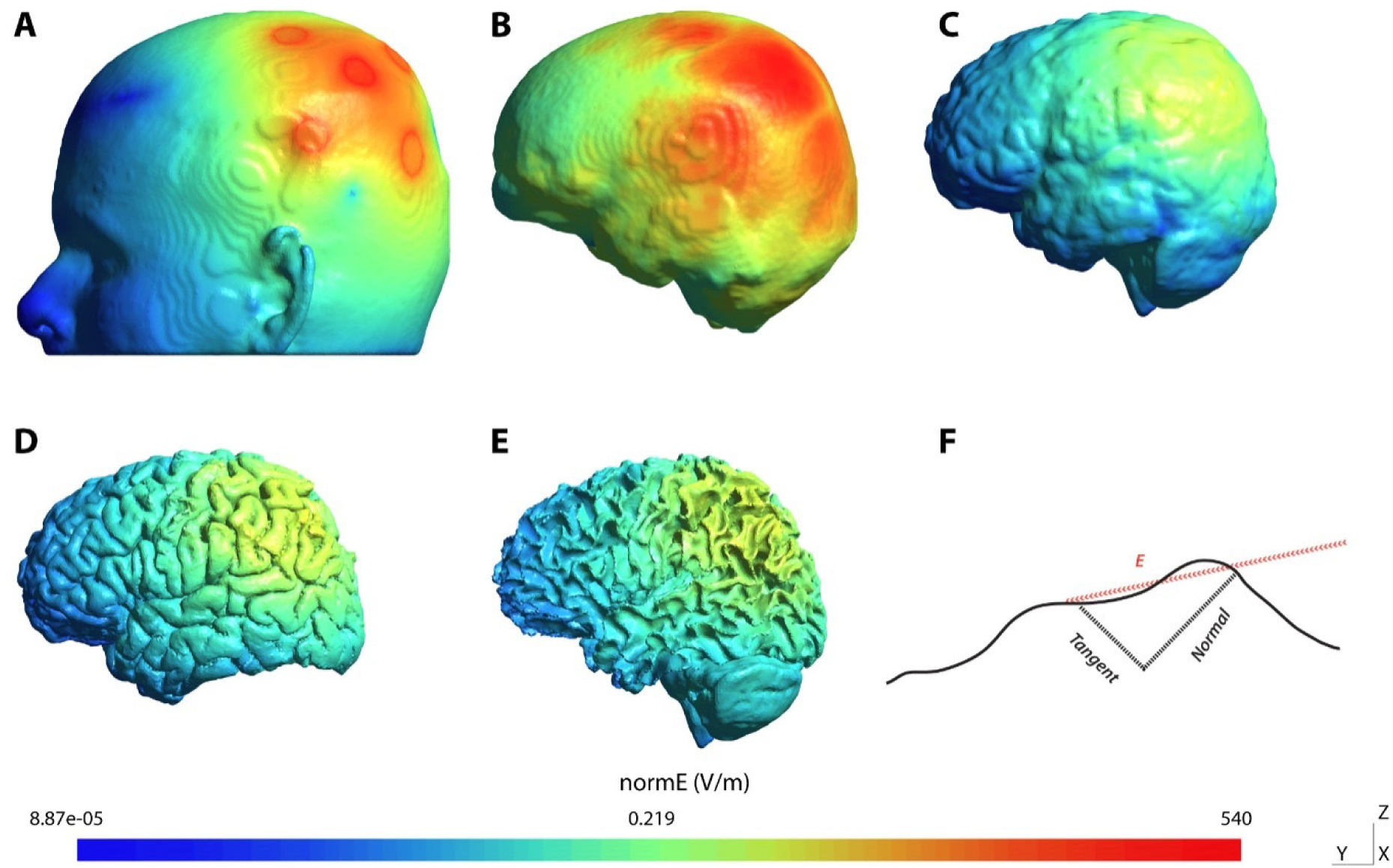
Source reconstruction of HD-tDCS over mIPS. The left medial intraparietal sulcus (mIPS) was localized during a separate functional MRI scan (Methods: Localizer task), and superimposed over a segmented head model (A-E). T1-weighted structural scans were segmented into tissue categories for finite element modeling of the electric field induced by HD-tDCS (Methods: Current Forward Modeling). Visualized is the electric field density across the scalp (A), skull (B), pial layer (C), gray matter (D), and white matter (E), showing a progressive decrease in the strength of the resulting electric field closer to neural tissue. Current models were performed in SimNIBS, with tissue conductivity values given in Table 1.

The MR images were segmented into tissue categories (scalp, skull, cerebrospinal fluid, gray matter, white matter) using an automated procedure in Freesurfer (Dale et al, 1999). This generated a tetrahedral mesh that was used as the model for simulating volume conduction in SimNIBS (Saturnino et al., 2018). Tissue conductance values used are reported in Table 1. The localization of mIPS was transformed from the Brainvoyager System Coordinates to SimNIBS space to guide the central electrode placement in the model with a custom Matlab function. Electrodes are modelled as elliptical nodes of dimensions 1cm x 1cm x 1mm on the tetrahedral surface. A ring mesh was generated with a 3cm radius to automatically place the 4 surrounding electrodes. Center electrodes were set to apply ± 2mA, and each surrounding electrode -/+ 0.5mA. The system of partial differential electromagnetic equations was solved using a conjugate gradient method to compute the vector field.

**Table 1:**
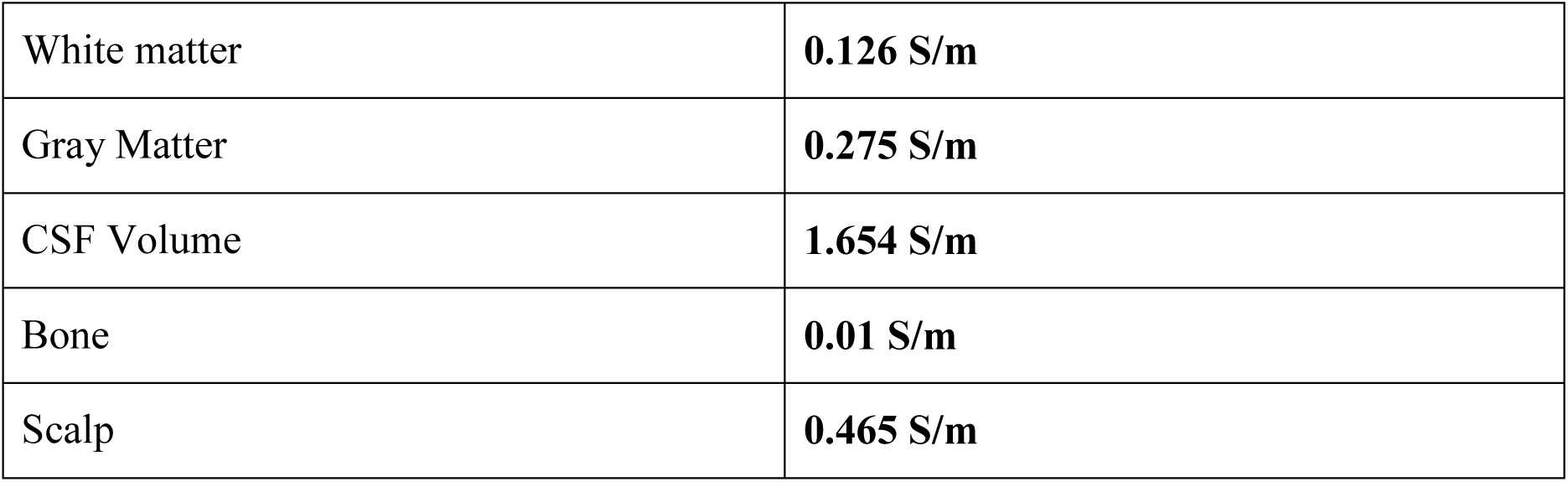
Tissue Conductivity Values based on Wagner et al., 2004.

|  |  |
| --- | --- |
| White matter | **0.126 S/m** |
| Gray Matter | **0.275 S/m** |
| CSF Volume | **1.654 S/m** |
| Bone | **0.01 S/m** |
| Scalp | **0.465 S/m** |

The resulting electric field is a vector field with a magnitude and orientation at the target site. We thus computed the magnitude of the radial and tangential (orthogonal complement of radial) component of the induced current at the localized mIPS site for a regression analysis with EEG and behavior. Electric fields were visualized with Gmsh (Geuzaine & Remacle, 2009) and MATLAB.

### EEG Analysis

We use the EEG analysis to test if changes in brain activity are directly related to changes in the online correction behavior. Of special interest, are particular brain oscillations that are thought to be a mechanism for cortical communication during motor planning and execution (Rubino, Robbins, & Hatsopoulos, 2006; see Battaglia Mayer, Ferrari-Roniolo & Visco-Comandini, 2015). Specifically, osciallations in the beta-band (15-30 Hz) have been implicated in the communication between the PPC and premotor cortex (Pani et al. 2014; Stetson & Anderson, 2014), somatosensory cortex (Zhang et al. 2008; Tsujimoto et al., 2009) and primary motor cortex (Menzer et al., 2014). In addition, the beta-band power is thought to be inversely related with movement preparation (Pani et al. 2014; Stetson & Anderson, 2014) which is crucial especially for online correction. To quantify the power of the oscillations, data was band-pass filtered between 1 and 45 Hz using a zero-phase digital filter. A notch filter excluded frequencies between 55 and 65 Hz to filter noise from the power line frequency. Each electrode was filtered independently. Trials were then aligned in separate analyses to the hand movement onset. A surface Laplacian was computed with respect to the central electrode in order to topologically localize the filtered data to the point on the scalp closest to mIPS (Babiloni et al, 2001). As we were interested in the online correction performance we used only the trials where a target jump occurred and due to the lateralization of the stimulation we also separated the analysis for trials jumping to the left or right.

For the time frequency analysis, data was analyzed in a 750 ms window covering [-250,500] ms relative to the target jump. Time-frequency energy representations were computed using a complex Morlet wavelet (BrainWave Toolbox; Jobst et al., 2018) with a width parameter of 7 centered on each time (0-500ms) and frequency (1-45 Hz) following the method of Tallon-Baudry et al (1997). The data was baseline variance-normalized using the z score with respect to each time vector. We computed such a representation for each baseline and post-stimulation session for the anodal and cathodal stimulation and again separated targets jumping to the left and right. Then we again quantified the effect of the stimulation by subtracting the baseline representation from that in the post-stimulation session.

### Mediation analysis

To examine the relationship between the behavioral changes, the modeled current flow, and the EEG response, we performed a mediation analysis (Baron & Kenny, 1986; see MacKinnon et al., 2007). Due to the lateralization of the stimulation, this was completed separately for trials jumping to the left and to the right. First, we established a relationship between the induced current based on the forward modeling and the behavioral changes. In a second step we correlated the induced current with the changes in the EEG to establish a relationship between stimulation and the potential mediator. To do so, we correlated the current vector with each point in the time frequency representation (TFR). By doing so we found two spots in the TFR response that were correlated with the induced current (first 100 ms after target jump for 11to 14 Hz and 100 to 200 ms from 20 to 26 Hz). To be able to represent that correlation and use it for the mediation analysis, we averaged the TFR differences in these time windows and took it as a measure of induced changes in the EEG. Following this, we correlated the changes in EEG with the behavioral changes. After these individual steps, the crucial step was now to compute the partial correlation between the induced current and behavior while controlling for the changes in EEG, allowing us to establish the causal structure of our stimulation.

In summary, to establish a causal relationship between stimulation-induced behavioral changes, mediation analysis should show three results: (1) correlation between stimulation current and behavioral change; (2) stimulation current changes brain activity as reflected in EEG rhythms; and (3) altered brain activity (change in EEG rhythms) determines behavioral changes.

### Simulating mediation statistics

In order to strengthen the interpretation of the results of our mediation analysis, we performed post-hoc Monte-Carlo simulations of our analysis pipeline for different true effect sizes and different numbers of participants. In order to generate two random variables *x*_*1*_ and *x*_*2*_ that are correlated with effect size *eff*, we used the following expression:

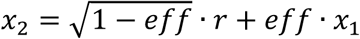

where r∼𝒩(0,1) is a normally distributed random variable with 0 mean and unit variance. As a result, both *x*_*1*_ and *x*_*2*_ are also normally distributed. Note, that for regressions the effect size *eff* is identical to the absolute value of the Pearson correlation.

We first generated a random variable (𝒩(0,1)) to represent the stimulation current. Using the above expression, we then generated the change in EEG signal and used this change in EEG signal to generate the change in a behavioral variable. We then ran n=5000 iterations for each effect size and number of participants combination, where we computed the significance of the relevant correlations and partial correlations between variables matching the three criteria outlined in the mediation analysis (see previous section). Specifically, for true positives, we required (1) significant behavior-current correlation, and (2) significant partial correlation between EEG and behavior, but (3) no partial correlation (insignificant)between current and behavior when accounted for EEG (significance threshold = 0.05). We did not require a significant current-EEG relationship because this analysis was done for each time point and frequency band and thus is likely to provide at least some significant result due to multiple comparisons. However, if this were the case, we would not expect that the current-EEG correlation is predictive of behavior. For the true negative rate, we required no current-behavior and no EEG-behavior correlation. The results of this mediation statistics simulation are shown in Figure 3 for different effect sizes as a function of number of participants (sample size). When there was no real effect (*eff* = 0), the (false) positive rate was <0.0005 for all sample sizes. This is very encouraging; it essentially means that if we find a significant effect, we can be confident that the effect is real. As we will show below, all our reported significant effect sizes are >0.6. For our sample size, we found that the true positive rate was rather modest (<0.2) and the false negative rate remained high (>0.15). Thus, we cannot confidently interpret non-significant results as an absence of evidence.

**Figure 3.**
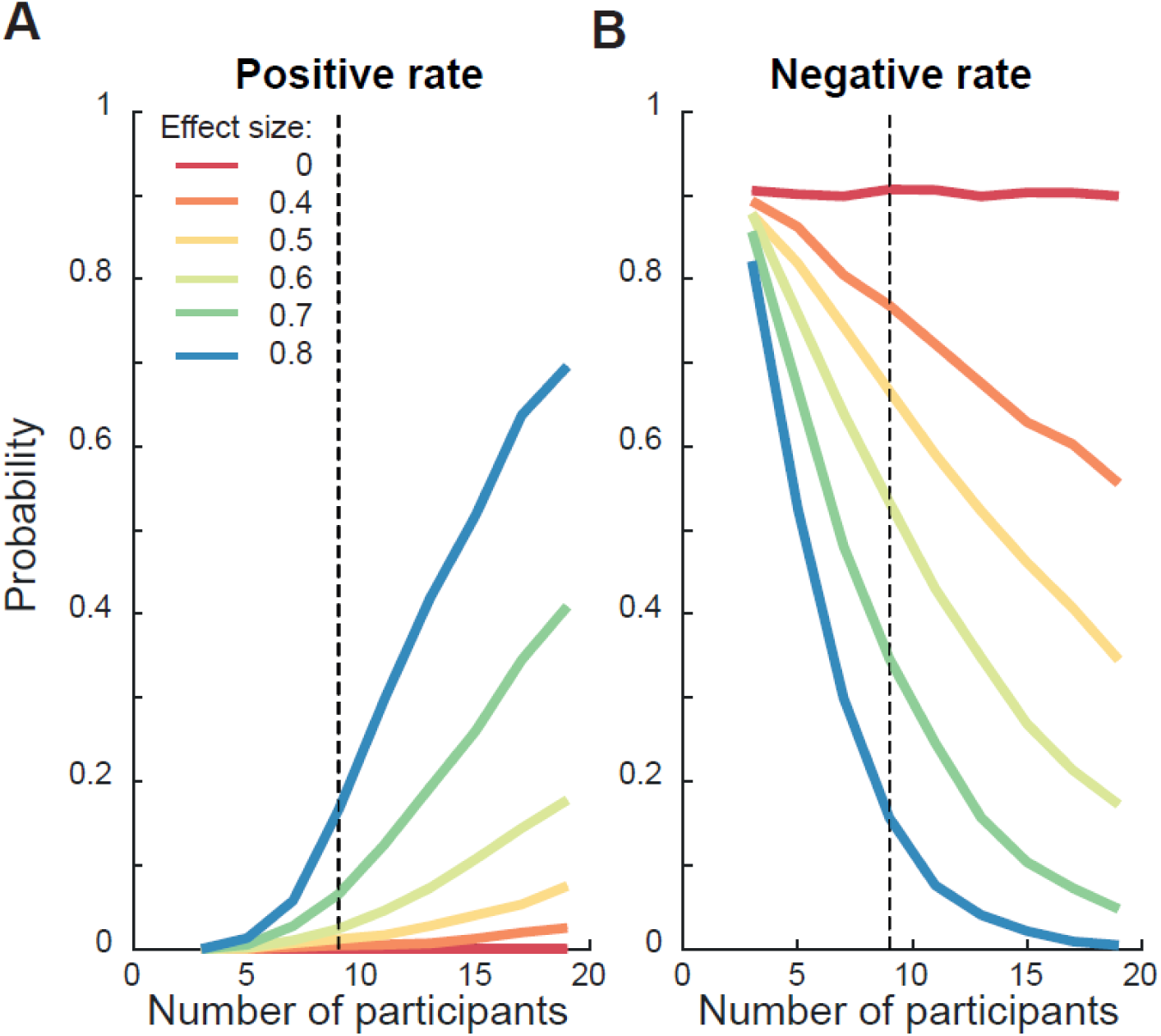
Simulation of mediation statistics. Depicted is the simulated probability of the (false) positive rate (**A**) and negative rate (**B**)given different numbers of participants (on the horizontal axes) and varying effect sizes (in different colors). The dashed vertical line represents the number of participants in the present study.

### Exclusion of trials

We had to exclude some trials from our analysis due to a missing signal of the Optotrak marker or where we could not determine hand movement onset or offset (∼6.5% of trials). Additionally, we excluded trials with hand movement onset latency faster than 100 ms or slower than 700 ms, indicating that participants probably did not conform tothe task (∼5.9% of trials). For the online correction targets, we excluded trials where the maximum deviation between the trajectory to the central target and the jumped target was less than 3 cm and the calculated online correction latency was outside of 100 and 700 ms, indicating that no appropriate correction had been made (<0.1% of trials). To calculate the average value for the baseline session we excluded trials from the first block, in order to eliminate contamination from any practice effects. This led to a total of 13850 out of 15840 trials (9 Participants *1920 trials - 9 Participants * 2 Days* 80 trials for the exclusion of the first block). For our EEG analysis we had to exclude the data from one participant as due to technical problems with determining the timing of the trials in the EEG-recording.

Data and analysis code for this project is available at: https://osf.io/kbyxc/

## Results

We used HD-tDCS over the left mIPS to elucidate the causal role of the PPC for online-control during reaching movements. In our task, participants had to perform reach-to-point movements to three different target locations. Occasionally the central targets would jump to the left or right at hand movement onset, thereby eliciting an online adjustment of the movement trajectory. We hypothesized that, depending on the polarity of the stimulation, there would be a differential effect on the online correction: cathodal stimulation, which should lead to a reduction of activity similar to artificial or real lesions, should lead to worse online correction performance. This would be reflected in a higher curvature of the movement. Anodal stimulation, which should lead to an increase in activity, should lead to an improved online correction performance, thus a lower curvature of the movement. In a next step, we used two independent measures of the changes in brain activity, one based on reconstruction of the current flow and one based on changes in EEG activity, and related these to the behavioral changes. The combination of our measurements allowed us to unravel the causal structure of the stimulation effect by linking them in a mediation analysis.

## Behavioral Analysis

### Movement accuracy and precision

To verify that participants were able to adjust their movements online we examined the trajectories of the individual arm movements (see Figure 4A). Participants were able to correct their movement in flight and land close to the same position as for the static movements. In order to assess the performance of the reaches we analyzed at the endpoints of the reaching movements. We calculated the horizontal- and vertical-position error as measurements of the accuracy and fitted an error ellipse to the endpoints of each condition as an indicator of the precision (see Figure 4B). We did not observe any systematic modulations of the horizontal error or precision of the movements and the endpoints for the corrected movements only had a higher vertical error than the comparable static movements across the different sessions (F(1,8) = 15.061, p = .005 for jumping left; F(1,8) = 11.900, p = .009 for jumping right). Importantly, there was no polarity specific effect on the endpoints for any of the measures. Based on this result, we continued with analyzing online control of the reaching movement with confidence that any polarity-specific effect we observe there cannot be explained by a systematic variation of the endpoint based on the stimulation.

**Figure 4.**
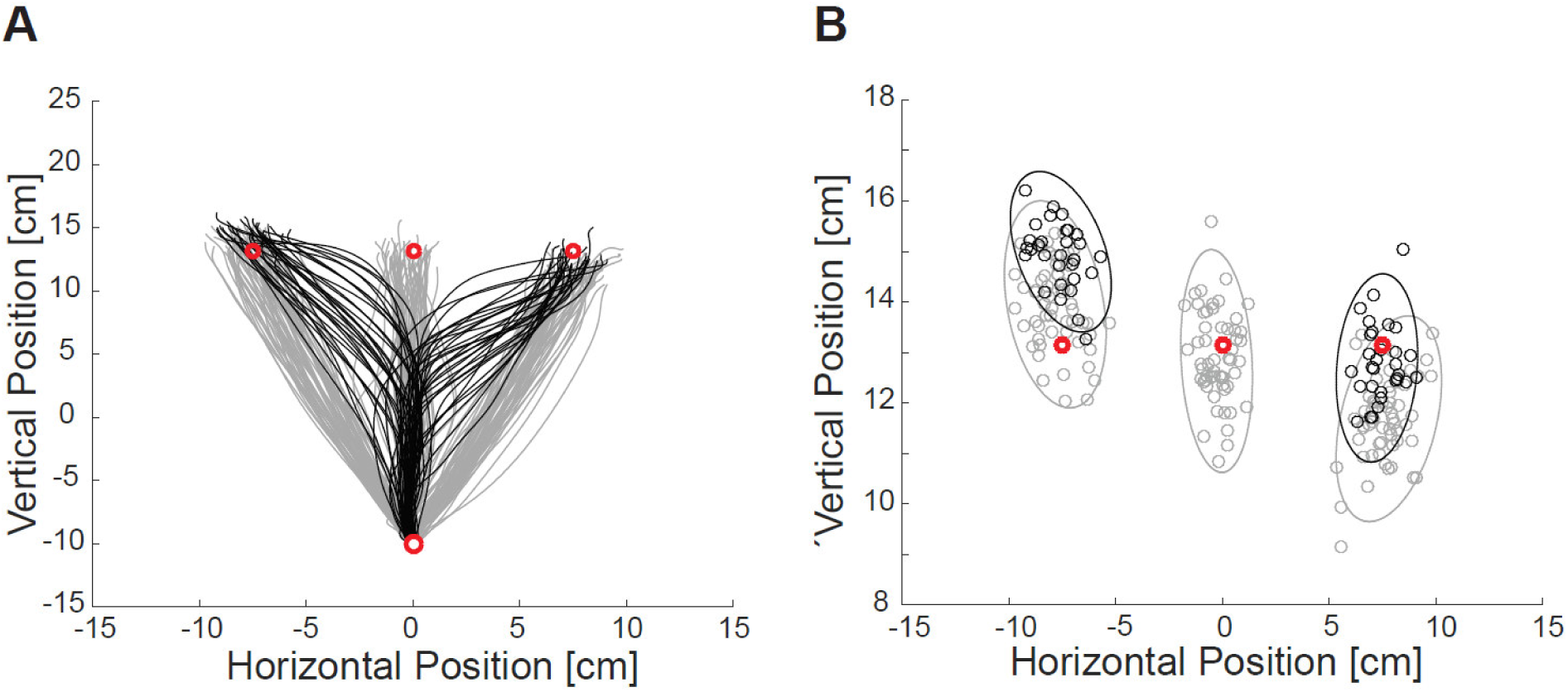
Depiction of hand movements. (**A**) Example reach trajectories of one baseline session for one representative participant. Grey paths show movements to static targets, black lines show movements to targets that jumped at hand movement onset. The red circles show the starting position and the possible target locations. (**B**) Corresponding end points of the reaches in (A). Again grey circles show the end points of movements to static targets, the black circles for the targets that jumped at hand movement onset. The red circles depict the target location. Separate error ellipses were fitted to each of the possible conditions.

### Movement curvature

To measure the effectiveness of the online correction we looked at the normalized curvature of the reaching trajectories. As mentioned earlier, a lower curvature indicates a smoother integration of the updated movement plan. A hypothetical maximal curvature value would be obtained if the movement was not corrected inflight but if the trajectory would be based on two completely separate movements. We used a repeated measurement ANOVA with three factors (polarity, target location, session) to test for possible effects on the online correction behavior. We found a significant main effect of session (F (2,16) = 4.584, p = .027, η^2^ = .364) and two interactions. The main effect was driven by a decrease in curvature across each session (Base: M = 0.160 ± 0.031; Stimulation: M = 0.154 ± 0.030; Post-Stim: M = 0.149 ± 0.028), suggesting that participants generally improved during the course of the experiment.

The interactions showed that there was an additional effect caused by the type of stimulation. We found a significant interaction between polarity and session (F (2,16) = 5.216, p = .018, η^2^ = .395) and between target location and polarity (F (1,8) = 6.514, p = .034, η^2^ = .449). While the curvature was slightly higher for the anodal stimulation blocks during the baseline, there was a reduction in curvature during anodal stimulation, which did not occur with cathodal stimulation (see Figure 5A). We investigated this effect further by running two repeated measures ANOVAs only with the factor Session for the anodal and the cathodal values, respectively. While we found a significant reduction over the sessions for the anodal stimulation (F (2,16) = 8.002, p = .004, η^2^ = .500), there was no reduction of the curvature for the cathodal stimulation (F (2,16) = 0.474, p = .631, η^2^ = .056). The interaction between target location and polarity was based on a higher curvature for cathodal than for anodal stimulation for rightward (contralateral) targets (see Figure 5B). There is a marginal reduction of the curvature for anodal stimulation for targets corrected to the right in comparison to the left (t (8) = 2.16, p = .063), but no difference depending on the direction with cathodal stimulation (t (8) = 0.16, p = .878). Thus, taken together there seems to be a reduction of curvature with anodal stimulation, whereas an increase in curvature with cathodal stimulation. This effect was most prominent for targets jumping to the right, i.e., the contralateral side of the stimulation.

**Figure 5.**
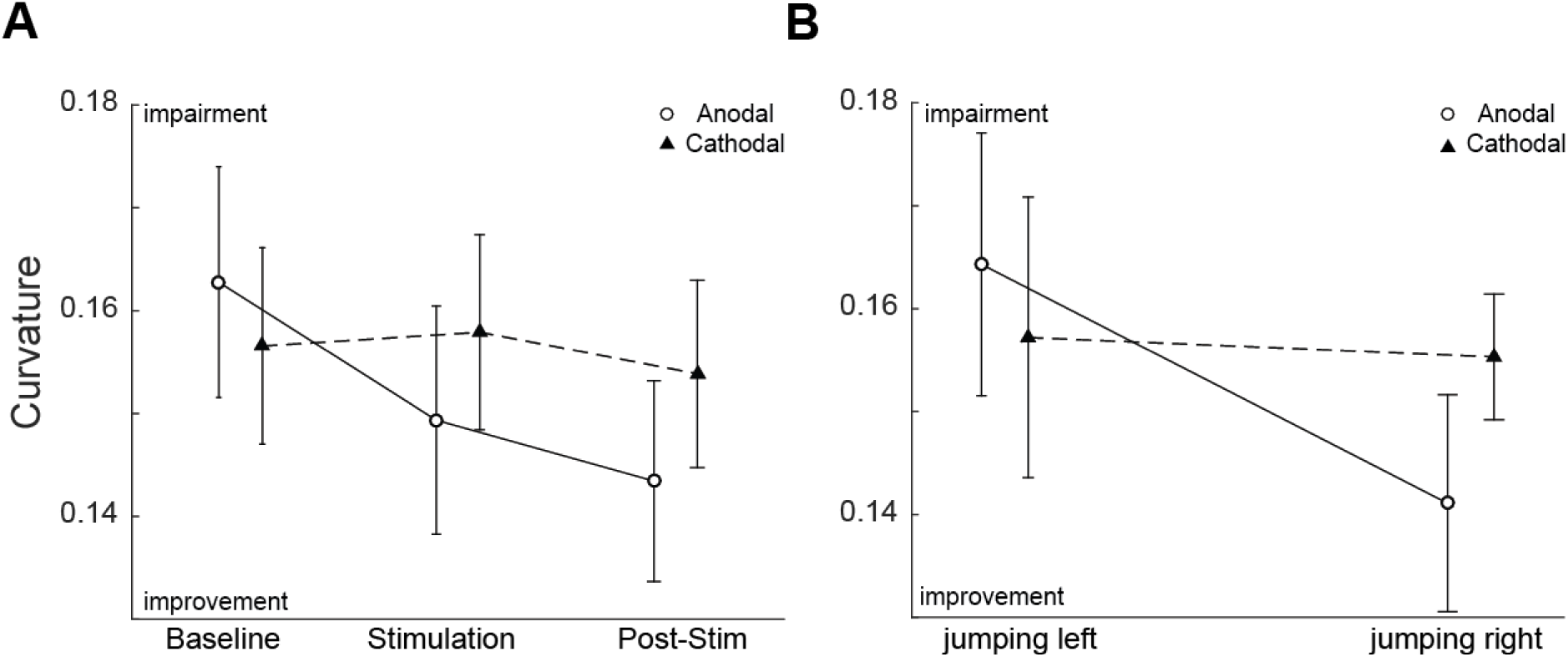
Effects on the curvature of the trajectories. (**A**) Curvature of the hand movements averaged across participants and shown separated for the different sessions and polarities (circle = anodal stimulation, triangle = cathodal). (**B**) Average curvature across participants for targets jumping to the left or right for both polarities. Error bars depict the standard error of the mean.

To investigate the integration of the new target location into the movement we also examined the development of curvature over time (see Figure 6A). We aligned all trials to the calculated onset of the online correction (see Methods) and calculated the curvature with a sliding window of 100 ms in steps of 10 ms. Interestingly, the latency of the online correction also decreased across the different sessions (F (2,16) = 10.093, p = .001, η^2^ = .558:Base: M = 291.70 ±63.42 ms; Stim: M = 280.43±64.54 ms; Post-Stim: M = 274.13±57.55 ms), but it was not affected by the polarity of the stimulation (F(1,8) = 2.539, p = .150, η^2^ = .241). To quantify the possible effect of the more effective integration of the updated movement plan, we normalized the curvature over time for each participant based on thei rpeak of the time course in the baseline condition and computed the difference between the peak curvature for the stimulation session and the baseline value. We did so for each of the polarities as well as for targets both jumping to the right and left (both polarities are shown in Figure 6A, but note that only trials jumping to the right are shown). We found a significant difference between the anodal and cathodal stimulation only during the stimulation for targets jumping to the right (t (8) = 3.725, p = .006), which was not present on the left side (t (8) = 1.37, p = .21). The fact that the contralateral side is significant but the ipsilateral is not, is not enough to show that both sides are actually different. Thus, we compared the difference between anodal and cathodal stimulation on both sides directly (Figure 6B) and again obtained a significant result (t(8) = 2.687, p = .028), indicating a hemifield specific effect of the stimulation. Furthermore, anodal stimulation led to a reduction in curvature, whereas cathodal stimulation led to an increase. Thus, in contrast to the latency of the online control system, we found a significant effect of the stimulation on the curvature of the reaching trajectories. Anodal stimulation seems to lead to a more effective updating and correction of the ongoing movement for targets that jumped into the contralateral field from the stimulation site, whereas the opposite was the case for cathodal stimulation.

**Figure 6.**
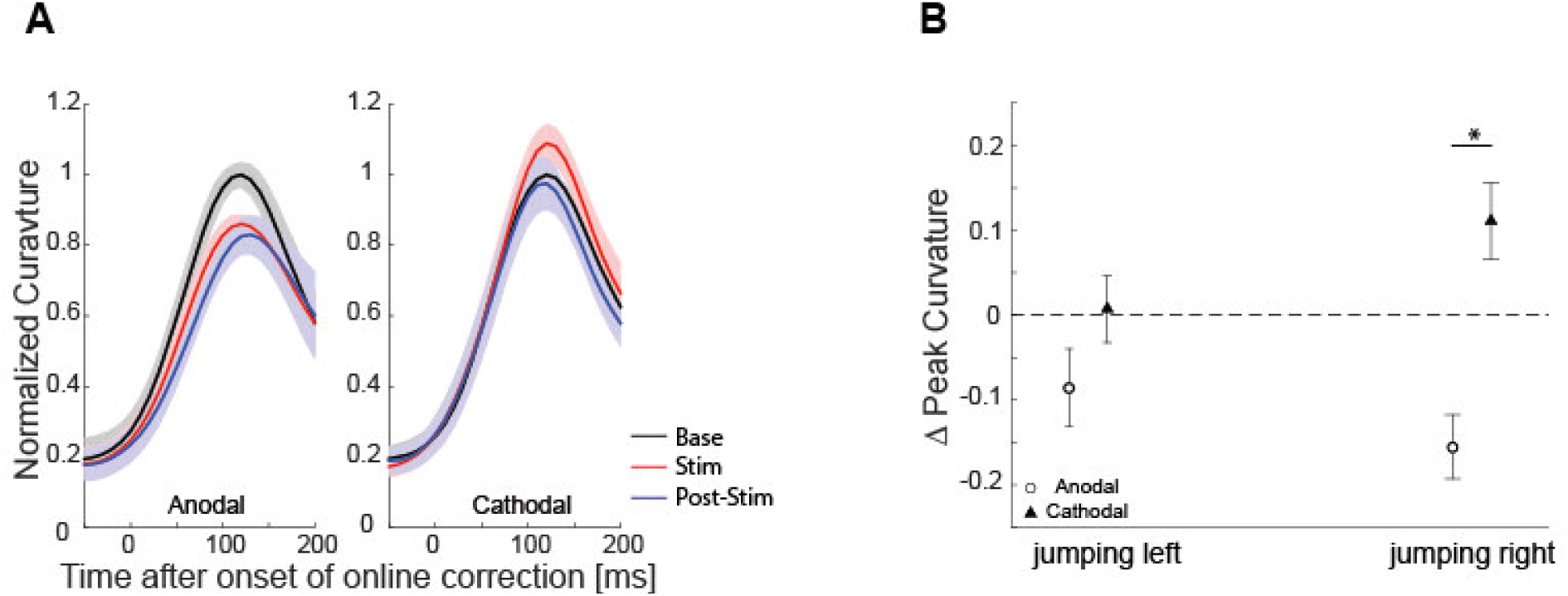
Development of curvature over time. (**A**) shows the average curvature over time aligned to online correction onset for the different sessions (Baseline = black, Stimulation = red, Post-Stim = blue). The left panel shows the curvature for targets jumping to the right during the sessions with anodal stimulation, the right panel with cathodal stimulation. Curvature was normalized so that the maximum of the baseline was assigned a 1. The shaded area shows the standard error of the mean. (**B**) depicts the average differences between the peak curvature during baseline and the stimulation. Circles show the effect of anodal stimulation, triangles of cathodal stimulation. Error bars depict the standard error of the mean. The star indicates significance at the 1% level.

### Causal relation between induced current and behavioral changes

Thus far our results indicate a relation between the polarity of the stimulation that we induced and participants’ online correction behavior. To test for a causal relationship, we next sought to provide evidence that the amount of current delivered to the mIPS, quantified based on an individual model of current flow for each participant (see current forward modeling in methods), is directly linked to their behavior and that this relationship is mediated by a change in brain activity. To establish such a relationship our data need to fulfill the following three criteria: (1) There needs to be a correlation between the induced current and the change in behavior. (2) There needs to be a relationship between the induced current and changes in brain activity, quantified in our case via EEG measurement before and after the stimulation. (3) The altered brain activity needs to be related to the changes in behavior and crucially this relationship must be the determining factor for the correlation between the induced current and the behavior.

### (1) Relationship between induced current and behavior

To test this relationship, we first quantified the induced current. We estimated the current that affected the left mIPS based on the electrode placement and the structural MRI scan of each participant (see current forward modeling in methods). The current was decomposed in its radial and tangential component. By design, anodal stimulation led to positive values, whereas cathodal stimulation led to negative values. We correlated the reconstructed current vectors with the effect of the stimulation on curvature and online curvature (stimulation-baseline for both measures). While there was no systematic relationship between the behavior and the radial component of the current (see Fig 7A), we observed a significant relationship between the tangential component of the current reconstruction and the curvature measurements for targets jumping to the right (see Figure 7B for online curvature). Importantly, and in line with the behavioral effects above this relationship was only present on the contralateral side of the stimulation, thus for targets jumping to the right.

**Figure 7.**
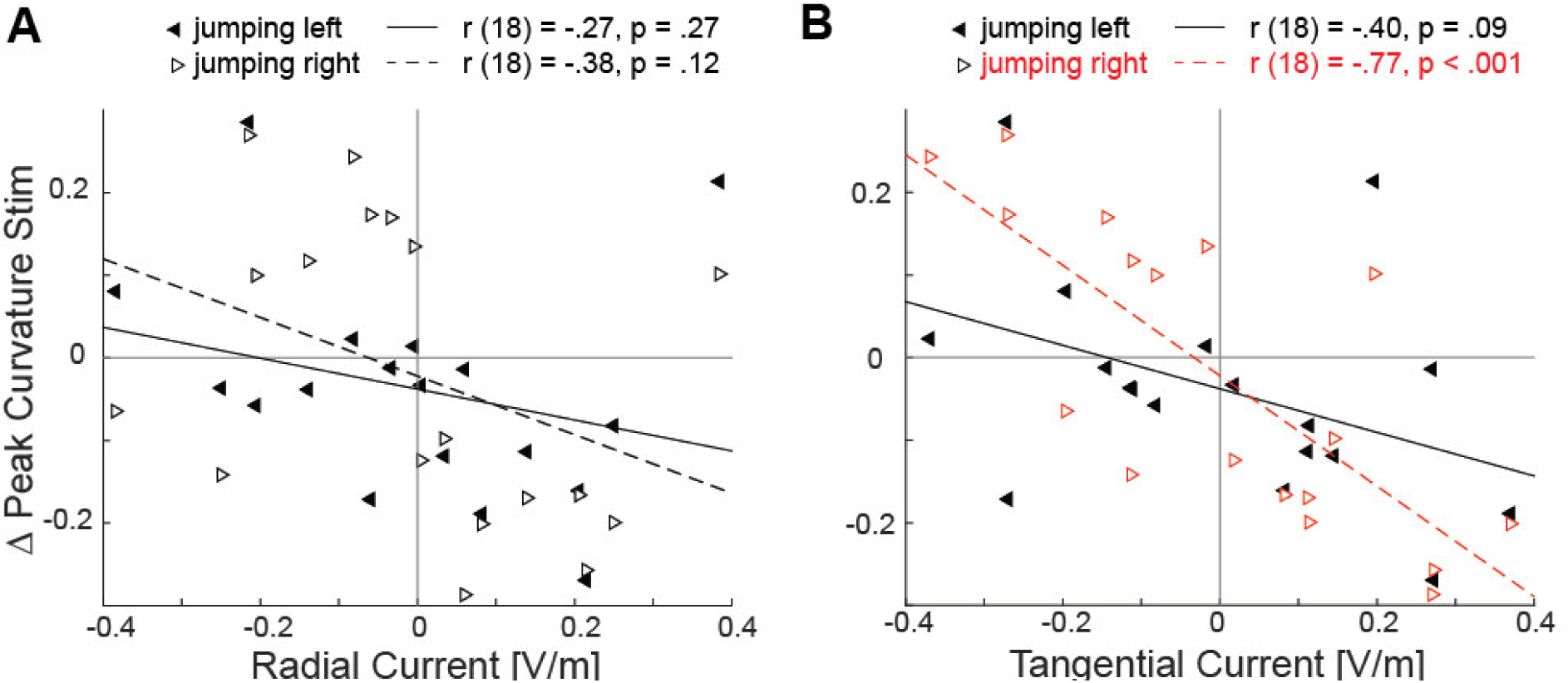
Relationship between induced current and behavior. **(A)** Relationship between the radial component of the induced current and the difference in peak reach curvature. Targets that jumped to the left are depicted by the black triangles pointing to the left. Targets jumping to the right are depicted by the open triangles pointing to the right. The solid line shows a regression for the targets jumping to the left, the dotted line shows the regression to the data jumping to the right. (**B)**Similar as A, but here the relationship with the tangential current is plotted. The red data points show the significant relationship between the tangential current and change in peak curvature for targets jumping to the right.

### (2) Relationship between current and EEG

We now have established the correlation between the induced current and the behavioral changes. To investigate whether the current also changed the brain activity, we analyzed the EEG response for trials that jumped either to the left or to the right. We measured the EEG in the baseline and the post-stimulation session, which allowed us to compute the effect of the stimulation on the brain activity. Recall that, during the stimulation, the sensors are being used for stimulation and thus cannot be used to record EEG activity. We computed TFR response for each session and direction of target jump separately and then computed the difference between the post-stimulation and baseline.

Figure 8 shows the difference in power across the different frequencies for anodal and cathodal stimulation respectively for targets jumping to the right. By looking at the figure it seems that anodal stimulation is slightly decreasing the power of alpha and beta frequencies shortly after the target jump, whereas cathodal stimulation seems to increase the power in a similar range (compare Figure 8A+B). To relate these changes with the induced current we wanted to establish areas in the TFR that were related to the induced current. Similar to the relation of the behavior, we found that the tangential current was related to the TFR responses for targets jumping to the right. Significant correlations were found quite localized at 11-14 Hz (alpha) in the first 100 ms after target jump and 20-26 Hz (beta) between 100 and 200 ms after target jump (see Figure 8C). As a representation of the observed correlation and in order toquantify the effect of the stimulation we extracted the average TFR response for targets jumping to the right for each participant in these time windows (see Figure 8D). For these time windows we found a significant difference between the effect of anodal and cathodal stimulation for targets jumping to the right (t (7) = 3.23, p = .015). Anodal stimulation led to a reduction of the power (M = -0.42, SD = 0.59), while cathodal stimulation increased the power (M = 0.22, SD = 0.28). This was true for the alpha as well as the beta activity, as the change in power in those two time windows was highly correlated (r (16) = .81, p <.001). If we computed the same difference for targets to the left we observed no significant correlation with the current and also no difference in the TFR representation (t (7) = 0.44, p = .67). Thus, we can establish a systematic influence of the induced current on the TFR response for targets jumping to the contralateral side of the stimulation. Note that this analysis was irrespective of any behavioral changes and only considered the lateralization of the brain with respect to coding of target jump direction.

**Figure 8.**
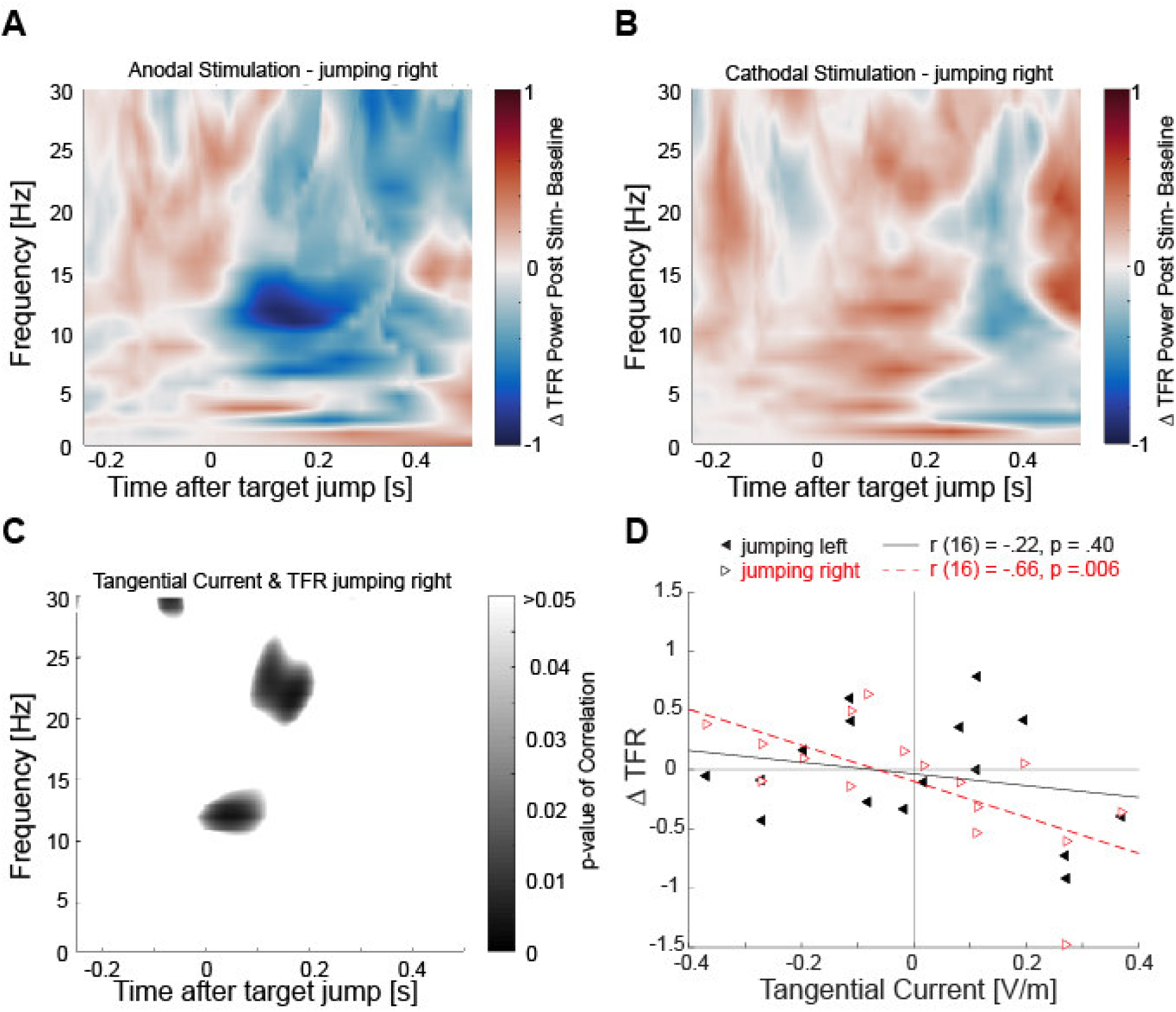
Relationship between induced current and EEG response. (**A+B)** Differences between Post-Stimulation session and Baseline in the TFR response aligned to the moment of the target jump. (**A)** shows the effect of anodal stimulation and (**B)** the effect of cathodal stimulation, both for targets jumping to the right. Blueish colors depict reduction of the power, reddish color increase in power. (**C)**depicts significant parts of a point-by-point correlation between the TFR response for targets jumping to the right and the tangential current. The p-value of these correlations is color-coded and to improve visibility only significant parts are shown. (**D)**Correlation between the average TFR values in the significant portions and the induced tangential current. Targets that jumped to the left are depicted by the black triangles pointing to the left. The significant relationship between tangential current and changes in the TFR for targets jumping to the right are depicted by the red open triangles pointing to the right. The solid line shows the regression for the targets jumping to the left, the dotted line shows the regression for the targets jumping to the right.

### (3) Mediation of the relationship between current and behavior via brain activity

Based on a causal interpretation of the effects, the induced current influences the brain response and this changed brain response should then affect the behavior. We now ask whether the independently established changes in EEG serve as a mediator for the relationship between current and behavior.

We tested for a relationship between the changes in EEG and behavior and observed a significant relationship between the changes in the TFR and curvature (r(16) = .62, p = .01; see Figure 9) as well as the curvature over time for targets jumping to the right (r(16) = .73, p =.001). Crucially, when we controlled for the changes in the TFR response by controlling for these changes in a partial correlation between the tangential current with the behavior, the correlations were significantly decreased (see Figure 8 for curvature; r (16) = -.54, p = .035 for online curvature).

**Figure 9.**
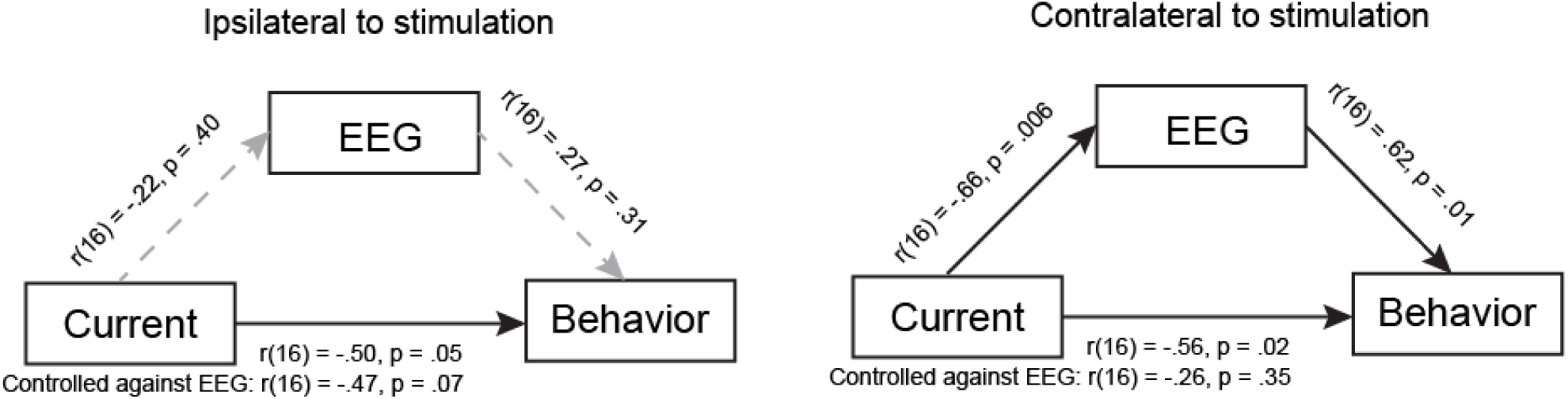
Mediation analysis. Relationship between the induced tangential current, the changes in the TFR response of the EEG and the change in curvature in the behavioral response. Solid arrows depict significant correlations. The partial correlation against the EEG response is reported beneath the relationship of induced current and behavior.

Thus, when taken together we found a polarity-specific modulation of the online correction for targets jumping to the contralateral side of the stimulation. Anodal stimulation slightly improved the correction, reflected in a lower curvature, whereas cathodal stimulation produced the opposite effect. We found a significant relationship between the tangential current and the observed behavioral changes and could successfully establish the changes in the EEG response as a mediator of this effect. These results suggest that the induced current influenced early alpha and beta oscillations after the target jump and these changes then affected the online correction performance. We only observed a successful mediation on the contralateral side, however based on our simulation of statistical power we cannot confidently rule out that the non-significant results for the mediation analysis on the ipsilateral side indicate the absence of an effect; it might also be possible that the stimulation tapped into a more general effect which affected both sides.

## Discussion

The goal of our study was to establish a causal role of the PPC for online reach control. We demonstrated not only behavioral changes, but more importantly, also reconstructed the causal structure of the effect of stimulation. We used a unique set of measurements: models of current flow for each individual participant, pre- and post-stimulation EEG recordings, and behavioral changes. Our results provide interesting insights into the role of the mIPS for online control, while establishing a new methodological approach to provide causality and explain some of the variability of tDCS stimulation.

### Online control behavior

Our results fit well into the literature on behavioral effects on online movement control (Gaveau et al., 2014) and optic ataxia (Rosetti et al., 2003; Andersen et al., 2014). Lesions in the PPC lead to severe impairments of online corrections (Grea et al. 2002; Buiatti et al., 2013), while keeping reaches to static targets intact (Desmurget et al., 1999; Grea et al., 2002). Here, we show that anodal stimulation, related to an increase in activity, leads to improved online correction performance, while cathodal stimulation, related to a decrease in activity, did not (Figure 5). Importantly, this effect was observed contralateral to the hemisphere of stimulation (Figure 6), providing evidence for the lateral specificity of stimulation. We did not observe any systematic changes in some more general behavioral measurements such as accuracy or precision. This finding rules out the possibility that our observed effects on online control are based on a change in the representation of the target position, suggesting that the stimulation indeed mainly affected the efficiency of online correction and not the endpoint of movement.

The exact behavioral and neural mechanisms of how movements can be adjusted in flight are still unknown; however recent optimal feedback control models provide a potential framework (Scott, 2004; Todorov & Jordan, 2002, Scott, 2012). These models propose that after establishing a behavior goal, the state of the body is estimated based on a combination of efferent and delayed afferent information. Motor actions are implemented by a control policy, which then continuously updates the movement (Scott, 2016; Medendorp & Heed, 2019). One key advantage of the optimal control theory framework is that it can explain why despite some variability in the individual trajectories (see Figure 4), the endpoints of a movement are still comparable and accurate.

In this framework the PPC seems to serve as a state estimator, due to its multiple afferent and connections and role in sensorimotor integration (Sirigu et al., 1996; Mulliken et al., 2008, Medendorp & Heed, 2019, Pilacinski & Lindner, 2019).Although our results provide additional evidence for the causal role of the PPC in online movement control, the PPC is not a uniform cortical area and different parts might fulfill different subtasks (Marigold et al., 2019, Medendorp & Heed, 2019). Our results suggest that our target area, the mIPS, is related to the integration of the incoming somatosensory information (Reichenbach et al., 2014). The stimulation seems to affect the internal state estimate, which is reflected in changes in the efficiency, but not the timing of online control.

One argument for a specific role of the mIPS in online control is that we only observed a significant mediation effect for targets on the contralateral side of the stimulation. However, since our simulation of the statistical power of the present approach revealed only a limited confidence in interpreting negative effects (Figure 3), it would also be possible that the effect of stimulating the mIPS is a more general one: instead of specifically affecting the integration of incoming information for targets on the contralateral side, it could have also broader effects on the integration of a new control policy.

### Brain oscillations as communication mechanism

Although the PPC is causally involved in online movement control, it is not theonly relevant area. Besides the PPC, the dorsal premotor cortex(PMd) plays a crucial role in updating target location (see Battaglia-Mayer et al., 2014). It provides a dynamic error signal indicating the new goal (Lee & Donkelaar, 2006), which then is passed to the PPC (Archambault et al., 2015). Interestingly, a lesion in the PMd mainly affects the latency of the movement, but not its accuracy (Buiatti et al., 2013). This is in line with our results, which show no effects on latency, but an impaired movement correction, presumably computed in the PPC (Battaglia-Mayer et al., 2014).

In order to adjust ongoing movements there are two crucial steps needed. First one needs to inhibit the old movement and afterwards integrate the state estimate of the hand with the new reach goal. These two steps seem to be related to specific frequencies of brain oscillations. The initial inhibition arriving from the PMd is presumably related to alpha oscillations (Pani et al., 2014), while the integration of the new goal and preparation of the movement may be related to beta frequencies (Engel & Fries, 2010; Stetson et al., 2014). Remarkably, we could reconstruct the two relevant frequency bands based on a correlation between our two independent measurements of changes in the brain activity, without taking into account the behavioral changes (Figure 8C). This correlation also provided a realistic temporal structure: an early alpha response after the target jump followed by a slightly later beta activity change.

The induced current influenced the power of alpha and beta frequency bands (Figure 8). Anodal stimulation significantly reduced alpha and beta power, suggesting that more efficient online correction is based on a combination of less inhibition of the old movement and an improved integration of the state estimate and the new goal, which is in line with above proposed hypothesis. Cathodal stimulation led to the opposite effect on both frequency bands and was related to a small increase in curvature. The change in both bands was highly correlated in our study (see Grabot et al., 2019), suggesting that despite their potential different functions, there might be a common underlying mechanism like the above described change in control policy (Scott, 2012).

### Caveats of results and strengths of themultimodal approach

Stimulation studies typically rely on the assumption that behavior won’t change in the absence of stimulation, but, trial-by-trial effects and learning in behavioral paradigms are the norm, not the exception (Fectau & Munoz, 2003). To statistically account for learning, manipulations of neural mechanisms should take ongoing changes in behavior into account. Our baseline/stimulation/post design allowed us to assess the improvement over time as well as the bidirectional effect of tDCS depending on the polarity (see Figure 5). Participants improved in the task across the different sessions, therefore a potential learning effect, in addition to a lack of statistical power, could hide additional effects. Based on the learning effect one possibility is that our results point to an increase or impairment of trial-by-trial learning, with anodal stimulation increasing learning and therefore leading to smoother corrections. To rule out that explanation, future work could add an additional sham conditionto measure the change in behavior without any stimulation. However, since we did not observe any polarity specific effect on other parameters like accuracy or latency of online correction and the observed changes were only present on the contralateral side of stimulation (with the caveat of not interpreting the negative effect for the ipsilateral side with too much confidence), our results point toward a direct influence of the mIPS for online reach control. Delivering current to humans noninvasively has been met with significant methodological challenges; much of the current delivered transcranially is shunted by the scalp, and the rest spreads diffusely across the cortex (Vöröslakos et al., 2018). Recent attempts to model the induced electric field has provided the opportunity to design electrode montages to increase the specificity of the induced current; like the HD-tDCS stimulation that we used in our experiment (Dmochowski et al., 2011; Saturnino, et al., 2015; Saturnino, et al., 2018). Additionally, the current flow depends on the individual anatomy (Miranda,et al., 2006), which may explain some of the variability and conflicting results in tDCS studies (Hill, et al., 2016; Kang et al., 2016). However note here, that not only the stimulation can be variable, but also the labeling of areas that people refer to as mIPS (Gallivan & Culham, 2015). Our setup allowed stimulation within a 3 cm radius, which is far more accurate than the length of the parts along the parietal sulcus, that are referred to as mIPS.

Despite these caveats, our experimental design contrasting anodal and cathodal stimulation combined with our multimodal approach (event-related EEG), fMRI-guided localization of stimulation target, and individualized current modeling work allow us to account for most of the potential problems. Specifically, modeled current source strength correlations with stimulation-induced EEG changes produced an independent subject-by-subject predictor for expected tDCS-induced behavioral changes. By independently measuring the change in brain activity via EEG, we could validate our current reconstructions due to the relationship between the induced current and changes in relevant frequency bands in the EEG. Based on the combination of our multiple measurements, it is possible to establish the causal structure of the stimulation while at the same time gain insight into potential mechanisms of online control.

Together, our design allows to integrate and make use of the variability that is typically found in the results of tDCS studies (Jacobsen et al., 2012) by the use of a regression-based mediation analysis, as a step to make tDCS a more useful tool for research and clinical applications. Future work can leverage our causal methodology to probe how multiple brain regions interact following brain stimulation, and the combination of our measurements can be adapted to almost any behavioral task and stimulation of different areas. This is an important step towards translating knowledge of the adaptive brain to our understanding of functional localization of behavior.

## Author contributions

**AG** Conception and Design of experiment, Performed Experiments, Analyzed behavioral and EEG data and performed the mediation analysis, Interpreted the results of the experiments, Prepared figures, Drafted manuscript, Edited and revised manuscript, Approved final version of the experiment.

**BC** Performed current reconstructions and EEG data analysis, Interpreted the results of the experiment, Prepared figures, Drafted manuscript, Edited and revised manuscript, Approved final version of the manuscript.

**JJ** Performed current reconstructions and EEG data analysis, Edited and revised manuscript, Approved final version of the experiment.

**SX** Performed experiments, Analyzed fMRI data, Prepared figures, Drafted parts of the method section, Edited and revised manuscript, Approved final version of the experiment.

**JG** Designed fMRI task and fMRI analysis, Edited and revised manuscript, Approved final version of the experiment.

**JD** Designed current reconstruction pipeline, Edited and revised manuscript, Approved final version of the experiment.

**KF** Conception and Design of experiment, Interpreted the results of the experiments, Edited and revised manuscript, Approved final version of the experiment.

**GB** Conception and Design of research, conception and design of analysis methods, Interpreted the results of the experiment, Edited and revised manuscript, Approved final version of the manuscript.

